# Population genetics of Himalayan langurs and its taxonomic implications

**DOI:** 10.1101/2024.09.04.611248

**Authors:** M.R Sharwary, K. Arekar, P. Karanth

**Author notes:** **Correspondence** Sharwary M. R., Undergraduate Programme, Indian Institute of Science, Bangalore - 560012, India.

## Abstract

Himalayan langurs (*Semnopithecus schistaceus*) are one of the most widely distributed colobine monkeys found in the Himalayas from Pakistan in the west to Bhutan in the east. Further, their distribution encompasses a wide range of elevation (from the foothills of the Himalayas to 4,270 m above sea level) and is interspersed with numerous deep river valleys. In this study, we investigate the role of riverine barriers and elevational gradients in shaping the population genetic structure in these langurs. Previous mitochondrial marker-based broad scale studies suggested limited role of river valleys in shaping the phylogeography of these langurs. Here we have utilized nuclear microsatellites and a more fine-scale sampling to further explore this issue. Fecal samples were non-invasively collected from two Indian Himalayan states Himachal Pradesh and Uttarakhand based on distribution records from past studies. A total of 7 microsatellite markers were genotyped for these samples. The data were subjected to various analyses, including Neighbor-joining tree, PCoA, AMOVA, STRUCTURE, and paired Mantel test. The results show an overall lack of population genetic structure and a much higher geneflow along elevational gradient than across river valleys. Significant isolation by distance was also observed. Additionally, our results do not support splitting the Himalayan langurs into multiple species/subspecies based on elevational gradient.

## Introduction

Himalayan langurs (*Semnopithecus schistaceus*) are colobine monkeys (subfamily Colobinae) widely distributed in the Himalayan region of India, Nepal, and parts of Pakistan and Bhutan (Blanford, 1888; Pocock, 1939). They are one of the few colobine monkeys occupying temperate climate regions and are distributed from the foothills of the Himalayas to 4,270 m above sea level (Bishop, 1979). The taxonomy of these langurs has been in flux for decades (see Arekar et al., 2021 and the references therein) with some authors such as Groves (2001) recognising three species in the Himalayan region namely *S. ajax, S. schistaceus*, and *S. hector*. Whereas others such as Hill (1939), consider the Himalayan langurs to be a single species *S. schistaceus* with multiple subspecies. The distributions of these species/subspecies are largely based on elevation. For example, in Groves (2001), *S. schistaceus* and *S. ajax* are distributed above 1500-2000 m whereas *S. hector* is distributed from the foothills to 1800 m. Similarly, in the case of other colobines from the Himalayan region such as capped langurs (Choudhary, 2014) and golden langurs (Wangchuk et al. 2003), subspecies distributions have been delimited based on elevation. Nevertheless, these studies have not tested if this apparent difference in their morphology is due to lack of geneflow – and therefore genetic differentiation between highland and lowland populations – or just clinal variation along an elevational gradient. In this regard, Bishop (1979) argued that Himalayan rivers cut deep through mountains (in a north-south direction) and gene flow is more likely to occur between lowland and highland populations rather than across river valleys. Further, Wallace (1854) observed that the distribution ranges of different species of Amazonian monkeys are shaped by rivers and hence they act as barriers to dispersal. In recent times, numerous studies have shown that rivers play an important role in shaping the population genetic structure of primates (Eriksson et al. 2004, Anthony et al. 2007, Nater et al. 2013, Fünfstück et al. 2014, Mitchell et al. 2015). Van Elst et al. (2023) studied how rivers, elevation, and paleoclimate interact to determine population genetic structure and differentiation in the Gerp’s mouse lemur.

With respect to the Himalayan langurs, Khanal et al. (2018) using mitochondrial genes examined the role of Himalayan rivers in shaping their population genetic structure. They found that certain large rivers of Nepal served as a barrier to gene flow beyond isolation by distance. They also found a low correlation between genetic variation and altitudinal gradient. Arekar et al. (2021) used an integrative approach (based on molecular, morphological, and ecological data) to resolve the taxonomy of Himalayan langurs. Their study indicated that the Himalayan population of *Semnopithecus* was distinct from *S. entellus* of the north Indian plains and therefore must be assigned to a distinct species *S. schistaceus* as suggested by Hill (1939). Additionally, their study did not support splitting this species into multiple subspecies as mitochondrial haplotypes assigned to different species (as per Groves 2001) did not form distinct clusters. Further, Arekar et al. (2022), expanded the sampling into the Indian Himalayan region (west of river Mahakali) that included many river valleys of western Himalayas (includes the Indian states of Himachal Pradesh, Uttarakhand and the union territory of Jammu and Kashmir). This study was also based on mitochondrial markers and encompassed the entire distribution range of Himalayan langurs. Their results showed that there was much sharing of haplotypes across river valleys in western Himalayas. Their statistical phylogeography analysis supported an east (Nepal) to west (western Himalaya) colonization of these langurs. Overall, these studies suggest that certain deep valleys in the Himalayas do serve as barrier, particularly in Nepal, whereas elevational gradient does not seem to prevent geneflow. However, the three genetic studies on the Himalayan langurs discussed above were based on mitochondrial markers and these results need to be confirmed using nuclear markers.

To bridge this gap, here we have undertaken a more fine-scale sampling of Himalayan langurs from two Indian states (Uttarakhand and Himachal Pradesh) in the western Himalayas. These samples were genotyped for seven microsatellite loci to better understand the role of elevation and deep valleys in shaping the population genetic structure of Himalayan langurs.

## Materials and Methods

### Sampling, microsatellite amplification and genotyping

A total of 176 fecal samples were collected non-invasively (without animal handling) from 46 locations in the Indian states of Himachal Pradesh and Uttarakhand (Fig. 1). Multiple samples were collected from each location. Host DNA was extracted from these samples using the protocol as in Arekar et al., 2021.

**figure 1.**
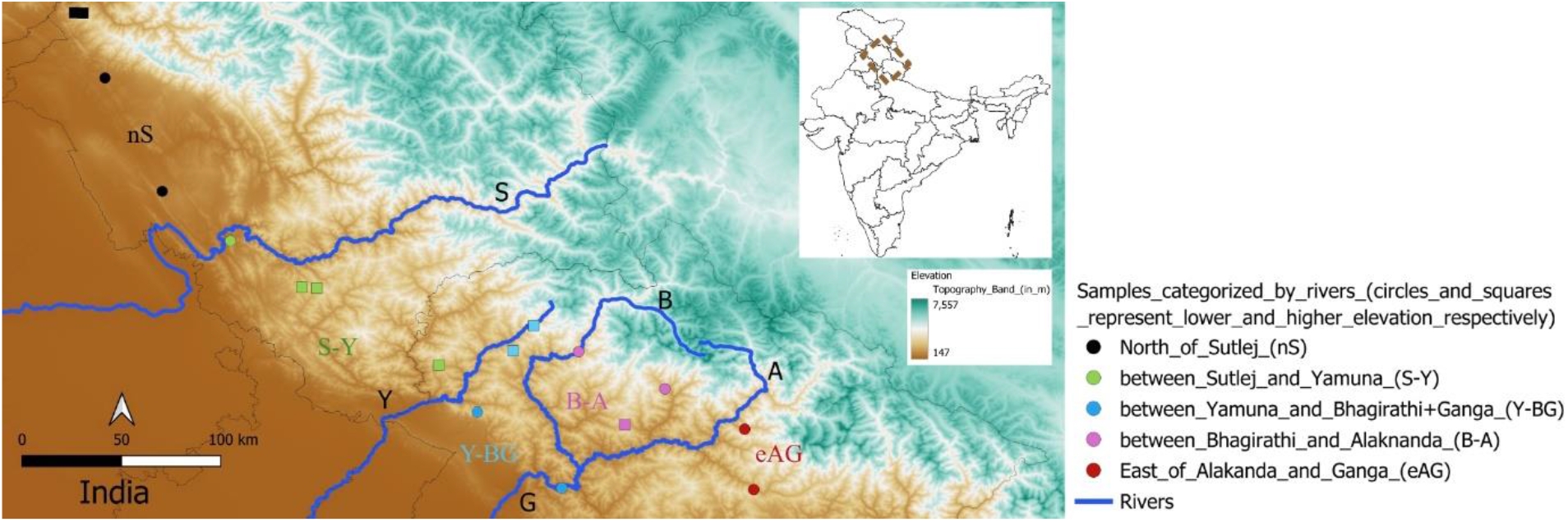
diagram showing the subpopulations of himalayan langurs with their label abbreviations separated by the five river valleys; s-river sutlej, y-river yamuna, g-river ganga, b-river bhagirathi, a-river alaknanda. the circles represent individuals from lower elevations and the squares represent individuals from higher elevations. the icons and labels of these populations correspond with those in nj tree (Fig. 2) and pcoa (Fig. 3).

The quality and concentration of extracted DNA were measured using a NanoDrop 2000 Spectrophotometer (Thermo Fisher Scientific Inc.). DNA concentration between samples usually varied from 20 to 80 ng/µl. The extracted DNA was aliquoted and diluted at a 1:5 (DNA extract: water) ratio to reduce PCR inhibitors during amplification. Due to the unavailability of published primers for microsatellite loci in Himalayan Langurs, 10 primers from the closely related genus *Trachypithecus leucocephalus* were used from Wang et al., 2016.

Out of the 10 selected microsatellite loci, 8 showed successful amplification during the preliminary trials. However, one of the primers was monomorphic. Hence, a total of 7 primers were finalized for the study (Table 1 given in Results section, refer to Wang et al., 2016 for specific details on the primers used). These microsatellite loci are fluorescently labeled (FAM dye) light-sensitive markers. Extra precautions were taken to minimize direct light exposure through the usage of aluminium foil and setting up reactions in low light conditions to prevent degradation of these primers.

PCR was set up using Ampliwin 2x PCR Master Mix with 0.03 µM concentration of primers, 2 µl of template DNA (4-16 ng/ul) in a 15 ul reaction volume. The thermocycler conditions were 95 °C initial temperature (5 min), 45 cycles of 95 °C denaturation (30 s), 49–54 °C annealing (45 s), 72 °C extension (45 s) and 72 °C final extension (10 min). Each primer was standardized through multiple temperature gradients separately. Table 1 (given in Results section) shows the annealing temperature of primers after standardization. Each sample was genotyped at least 3 times for each locus for consistent results and to reduce genotyping (allele calling) errors which are common when dealing with fecal DNA. Each PCR run had positive, and negative controls for amplification confirmation and quality control (checking for contamination).

The PCR products were outsourced to Barcode Bioscience, Bengaluru for genotyping, using LIZ500 as a size standard. The genotyping results were examined utilizing the microsatellite plug-in within Geneious Prime 2022.1.1 and the allele calling was done manually. The samples with inconsistent peaks or negligible amplification in three genotyping replicates were dropped. Individuals with missing genotypes for more than 3 of the 7 loci were removed from further analyses. Although we were able to collect 176 samples from our fieldwork, immediate DNA extraction and complete microsatellite amplification were challenging. Since it was non-invasive sampling, the extracted fecal DNA’s quality and concentration were low making it extremely tricky to amplify and ensure consistency in triplicate genotyping. We dropped several samples to maintain strong amplification and consistency in replicates to generate a robust genotypic dataset of 48 individuals.

The complete microsatellite dataset was used to calculate deviations from the Hardy-Weinberg Equilibrium and linkage equilibrium, using the Hardy-Weinberg Exact Test (Guo and Thompson, 1992) and G test respectively in Genepop v4.7.5 (Rousset, 2008). The program MICROCHECKER v2.2.3 (Van Oosterhout et al., 2004) was used to screen for genotyping artifacts such as the presence of null alleles, allelic dropout, and stutter products.

### Analysis of genetic diversity

We measured genetic diversity indices such as the total number of alleles, mean allelic richness, mean observed heterozygosity, and mean expected heterozygosity using FSTAT 2.9.4 and GenAlEx v6.5 (Peakall and Smouse 2006, 2012).

### Genetic structure and differentiation

Pairwise Nei’s genetic distance *D*_*A*_ (Nei et al., 1983) was calculated between individuals and used to construct the Neighbor-Joining (NJ) tree using Populations 1.2.31 (Langella, 1999) as well as Principal Coordinates Analysis (PCoA) to detect genetic clustering among individuals.

STRUCTURE v2.3.4 (Pritchard et al., 2000) software was used to detect any hidden population structure, irrespective of the geographical location of samples. It was run assuming a model of ancestral admixture and correlated frequencies for K values between one and ten, and 10 runs were performed for each K with 300,000 iterations following a burn-in of 50,000 steps. The optimum K was determined using STRUCTURE HARVESTER (Earl and vonHoldt, 2012) identify the real number of genetic clusters using the modal value of Δ*K*, a quantity based on the second-order rate of change of likelihood of the data (Evanno et al., 2005).

Isolation by distance was tested using the paired Mantel test wherein pairwise Wright’s Fst values were regressed on to the shortest geographic distances between sampling sites in GenAlEx v6.5 (Peakall and Smouse 2006, 2012).

To investigate the role of elevational gradient on genetic structure and differentiation, populations were assigned to putative species as per Groves (2001) classification scheme. To this end, samples that were collected between 600-1800m were assigned to *S*.*hector* and samples from elevational range of 2000-3000m were assgined to *S*.*ajax* (the two putative species distributed in the study area, refer to Fig. 1). Elevation data for sampling locations was extracted from the DEM database using latitude and longitude coordinates (GPS Visualizer). These putative species were subjected to Analysis of Molecular Variance (AMOVA) using Nei’s genetic distance in GenAlEx v6.5 (Peakall and Smouse 2012) which allows the hierarchical partitioning of genetic variation within and among populations. Similarly, to investigate the role of river valleys on genetic structure and differentiation, sampled populations were segregated into five subpopulations that were separated by rivers Sutlej, Yamuna, Bhagirathi and Alaknanda (refer to Fig. 1). These five subpopulations were subjected to AMOVA analysis. The program was run with 9999 permutations at 0.95 significance levels.

## Results

### Genotyping and analysis of genetic diversity

Most of the microsatellite loci (except WHL60) showed deviation from Hardy-Weinberg Equilibrium and signs consistent with null alleles. There was no evidence of large allelic dropout or stutter products in any of the seven loci. However, the Hardy-Weinberg exact test in Genepop is known to overestimate the evidence for homozygote excess (Engels 2009). Additionally, Dharmarajan et al., 2013 showed that currently employed quantitative methods like Microchecker cannot reliably distinguish between heterozygote deficits resulting from null alleles and those caused by biological mechanisms like Walhund effect. Hence, our analyses include the entire dataset since the deviation from HWE can be due to Walhund effect due to the pooling of different individuals/subpopulations which is common when analyzing natural populations (Waples, 2015). The linkage disequilibrium test showed signs of linkage between two pairs of microsatellite loci (WHl122 and WT173, WHL60 and WHL121). Analyses were run with and without WT173 and WHL121, however, the results were not different (reported in Table S1), hence we report the findings including all microsatellite loci. The genetic diversity estimates of each of the loci are as given in Table 1.

**Table 1.** Genetic diversity estimates and annealing temperature per locus. Abbreviations: *H*_*O*_: Mean observed heterozygosity, *H*_*E*_: Mean expected heterozygosity, N_A_: Mean number of alleles, AR: mean allelic richness.

| Sr. no | Primer name | Annealing Temperature (°C) | $H_O$ | $H_E$ | $N_A$ | $A_R$ |
| --- | --- | --- | --- | --- | --- | --- |
| 1 | WHL-60 | 49 | 0.374 | 0.456 | 2.800 | 3.974 |
| 2 | WHL-138 | 60.3 | 0.471 | 0.561 | 3.800 | 6.708 |
| 3 | WHL-122 | 57.6 | 0.510 | 0.698 | 5.000 | 8.769 |
| 4 | WHL-121 | 54 | 0.305 | 0.535 | 4.200 | 6 |
| 5 | WT-111 | 54 | 0.040 | 0.173 | 1.600 | 3 |
| 6 | WT-173 | 50.3 | 0.592 | 0.716 | 5.000 | 8.974 |
| 7 | WT-181 | 53.5 | 0.576 | 0.782 | 7.000 | 13.974 |

The mean observed heterozygosities ranged from 0.305 (WHL-121) to 0.592 (WT-173 and mean allelic richness ranged from 3 (WT-111) to 13.974 (WT-181).

### Genetic structure and differentiation

In the NJ tree, individuals do not cluster either based on elevation or river valleys (Fig. 2). Individuals from these two assigned categories are intermingled in the tree. In the PCoA plot the first axis explained 12.5% of the variation. The second axis explained 11.45% of the variation while the first three axes explained 34.03% of the variation. Here again there is no clustering of individuals based on either river valley or elevation (Fig. 3). However, the North of Sutlej (nS) and East of Alaknanda + Ganga (eAG), the most distant pair putative populations were separated in the first axis. STRUCTURE did not detect any patterns of hidden population structure, irrespective of geographical locations, which was in concordance with PCoA (bar plot in Fig. S1).

**Figure 2.**
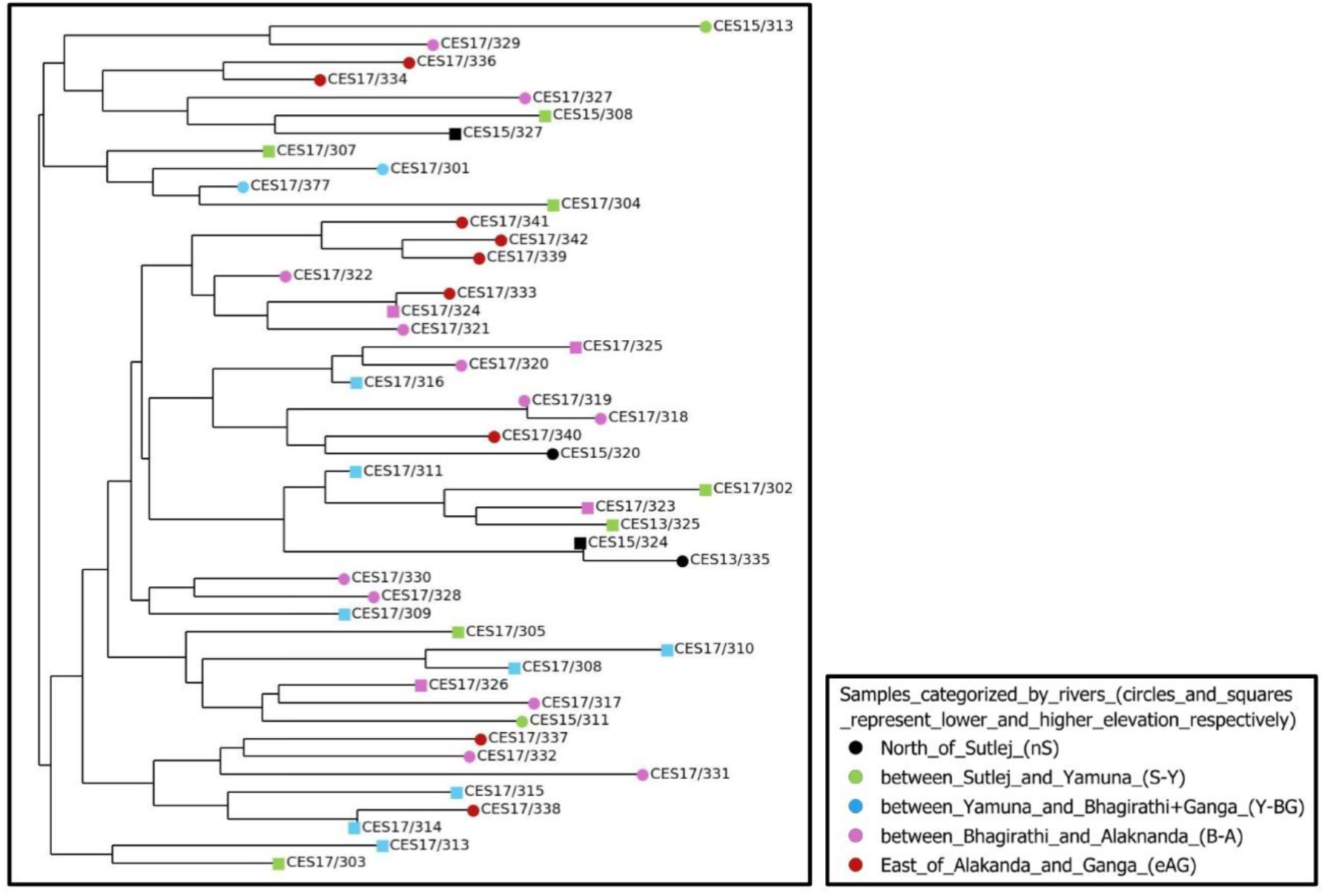
Neighbor joining (NJ) tree. The icons and labels correspond to those in Fig. 1 and Fig. 2.

**Figure 3.**
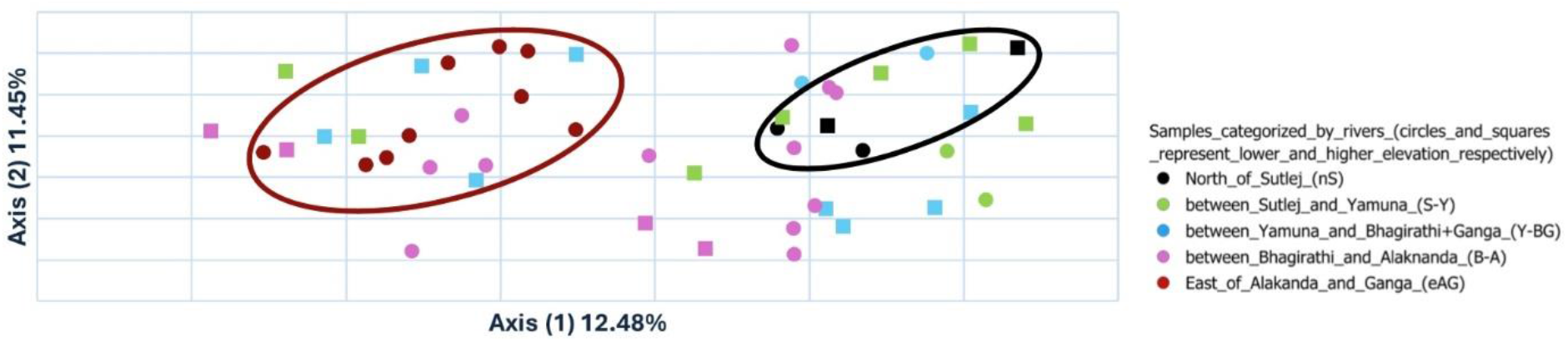
Principal Coordinate Analysis (PCoA). The larger circles, black and red correspond to the subpopulations nS and eAG respectively. The icons and labels correspond to those in Fig. 1 and Fig. 2.

In the analysis of molecular variance (AMOVA), when subpopulations were segregated based on elevation, 98% of the variation was distributed within the lower and the higher elevation subpopulations, and only 2% was contributed by variation between these subpopulations. However, this genetic differentiation between the lower and higher elevation subpopulations was not significant (AMOVA Φ_*PT*_= 0.023, P > 0.05). In the case of subpopulations separated by river valleys, 91% of the total variation was found within subpopulations and 9 % was distributed among subpopulations. The highest genetic differentiation was observed between the most distant pair of subpopulations, North of Sutlej (nS) and East of Alaknanda + Ganga (eAG) (AMOVA Φ_*PT*_= 0.333, P < 0.05), while genetic differentiation among most other subpopulations were not significant. The Mantel test showed a high positive correlation between the pairwise FST values (Y) between the subpopulations and shortest geographic distances (X) between them, indicating strong isolation by distance (Rxy = 0.823, P-value = 0.01, linear regression of XY plot in Fig. S2).

## Discussion

Riverine barriers, and to some extent elevational gradients, have been known to govern the population genetic structure in primates (Eriksson et al. 2004, Anthony et al. 2007, Nater et al. 2013, Fünfstück et al. 2014, Mitchell et al. 2015, Khanal et al., 2018, van Elst et al., 2023). However, there have been limited studies on this topic in South Asia, particularly with respect to testing the influence of elevational gradients on population genetic structure (Sekar and Karanth, 2013, Tyagi et al., 2016, Khanal et al., 2018). We focussed on Himalayan langurs, a colobine monkey with a history of convoluted taxonomy and wide distribution across the Himalayas making it a good model system to test the role of elevational gradients and riverine barriers on their population genetic structure. By using nuclear microsatellites, we attempted to get a better genetic resolution to elucidate their patterns of genetic structure and differentiation.

The NJ tree did not reveal any elevational or topographical clustering of the subpopulations. Similarly, PCoA and STRUCTURE analyses also did not retrieve any clustering of putative subpopulations. Nevertheless, in the PCoA analysis the most distant pair of subpopulations (North of Sutlej (nS) and East of Alaknanda + Ganga (eAG)) did separate out in the second axis. A mantel test confirmed significant isolation by distance, hence the genetic differentiation observed between the subpopulations can be attributed to the geographic distance rather than barriers associated with elevation or river valleys. Large rivers in the Himalayas are wide, perineal and fast flowing, making them formidable barriers for non-volant species. If so, how do we explain the overall lack of population genetic structure among Himalayan langurs in India? There are two possible explanations: First, animals might be able to cross further upstream where the rivers are narrow and have tree canopy on either side that meet over the river; Second, recently in the last ∼200 years, there are numerous suspension bridges that have been constructed across many of these rivers which might have facilitated animal movement (Bishop, 1979).

It must be noted that the AMOVA analysis suggested higher geneflow along elevational gradient as opposed to across river valleys (as suggested by Bishop, 1979). This result also calls to question Groves, 2001 classification scheme of Himalayan langurs, wherein the higher and lower elevation populations have been assigned to different species. The morphological difference between these populations is perhaps clinal variation in a single species along an elevational gradient (as noted by Choudhary and India, 2016).

Overall, our results suggested a limited role of river valleys in shaping population genetics of Himalayan langurs. Additionally, genetic differentiation among subpopulations of Himalayan langurs is largely governed by isolation by distance. These results do not support splitting the Himalayan langurs into multiple species/subspecies. We recommend subsuming all the known species/subspecies of Himalayan langurs into one species, *Semnopithecus schistaceus* (as in Arekar et al., 2021).

## Acknowledgments

We would like to thank the forest departments of Himachal Pradesh and Uttarakhand for providing the necessary permits. We would like to thank Ashika for her time to review microsatellite genotyping and Aritra for his help with review and support in the project. We would like to thank Dr. Theresa Burg, University of Lethbridge for suggestions on analyses.

## Author contributions

PK and KA obtained the funding for the project. PK and KA conceived ideas and designed methodology. KA collected the samples and SMR generated the data. SMR analyzed the data. SMR led the writing of the manuscript. All authors contributed to the draft and approved for publication.

## Supplementary Information

Additional information may be found in the online version of this article:

**Table S1.** Comparison of genetic diversity estimates across all loci and populations.

**Figure S1.** Bar plot of Structure.

**Figure S2.** Linear regression of XY plot from Mantel test.

## References

Anthony, N. M., Johnson-Bawe, M., Jeffery, K., Clifford, S. L., Abernethy, K. A., Tutin, C. E., Lahm, S. A., White, L. J. T., Utley, J. F., Wickings, E. J., & Bruford, M. W. (2007). The role of Pleistocene refugia and rivers in shaping gorilla genetic diversity in central Africa. Proceedings of the National Academy of Sciences, 104(51), 20432–20436.

Arekar, K., Sathyakumar, S., & Karanth, K. P. (2021). Integrative taxonomy confirms the species status of the Himalayan langurs, Semnopithecus schistaceus Hodgson, 1840. Journal of Zoological Systematics and Evolutionary Research, 59(2), 543–556.

Arekar, K., Tiwari, N., Sathyakumar, S., Khaleel, M., & Karanth, P. (2022). Geography vs. past climate: The drivers of population genetic structure of the Himalayan langur. BMC Ecology and Evolution, 22(1), 100.

Bishop, N. H. (1979). Himalayan langurs: Temperate colobines. Journal of Human Evolution, 8(2), 251–281.

Blanford, W. T. (1888). The fauna of British India, including Ceylon and Burma: Mammalia. Published under the Patronage of the Secretary of State for India.

Choudhury, A. (2014). Distribution and Current Status of the Capped Langur Trachypithecus pileatus in India, and a Review of Geographic Variation in its Subspecies. Primate Conservation, 28(1), 143–157.

Choudhury, A., & India, R. F. for N. in N. (2016). The Mammals of India: A Systematic & Cartographic Review. Gibbon Books and the Rhino Foundation for Nature in NE India.

Dharmarajan, G., Beatty, W. S., & Rhodes, O. E. (2013). Heterozygote deficiencies caused by a Wahlund effect: Dispelling unfounded expectations. The Journal of Wildlife Management, 77(2), 226–234.

Earl, D. A., & vonHoldt, B. M. (2012). STRUCTURE HARVESTER: A website and program for visualizing STRUCTURE output and implementing the Evanno method. Conservation Genetics

Engels, W. R. (2009). Exact Tests for Hardy–Weinberg Proportions. Genetics, 183(4), 1431– 1441.

Eriksson, J., Hohmann, G., Boesch, C., & Vigilant, L. (2004). Rivers influence the population genetic structure of bonobos (Pan paniscus). Molecular Ecology, 13(11), 3425–3435.

Evanno, G., Regnaut, S., & Goudet, J. (2005). Detecting the number of clusters of individuals using the software structure: A simulation study. Molecular Ecology, 14(8), 2611–2620.

FSTAT. (n.d.). https://www2.unil.ch/popgen/softwares/fstat.htm

Fünfstück, T., Arandjelovic, M., Morgan, D. B., Sanz, C., Breuer, T., Stokes, E. J., Reed, P., Olson, S. H., Cameron, K., Ondzie, A., Peeters, M., Kühl, H. S., Cipolletta, C., Todd, A., Masi, S., Doran-Sheehy, D. M., Bradley, B. J., & Vigilant, L. (2014). The genetic population structure of wild western lowland gorillas (Gorilla gorilla gorilla) living in continuous rain forest. American Journal of Primatology, 76(9), 868–878.

Geneious Prime 2022.1.1.(https://www.geneious.com)

GPS Visualizer: Assign DEM elevation data to coordinates. (n.d.). https://www.gpsvisualizer.com/elevation

Groves, C. (2001). Primate taxonomy. Smithsonian Institution Press.

Guo, S. W., & Thompson, E. A. (1992). Performing the Exact Test of Hardy-Weinberg Proportion for Multiple Alleles. Biometrics, 48(2), 361.

Hill, W. C. (1939). An annotated systematic list of the leaf-monkeys. Ceylon Journal of Science, 21, 277–305.

Khanal, L., Chalise, M. K., Wan, T., & Jiang, X. (2018). Riverine barrier effects on population genetic structure of the Hanuman langur (Semnopithecus entellus) in the Nepal Himalaya. BMC Evolutionary Biology, 18(1), 159.

Mitchell, M. W., Locatelli, S., Sesink Clee, P. R., Thomassen, H. A., & Gonder, M. K. (2015). Environmental variation and rivers govern the structure of chimpanzee genetic diversity in a biodiversity hotspot. BMC Evolutionary Biology, 15(1), 1.

Nater, A., Arora, N., Greminger, M. P., Van Schaik, C. P., Singleton, I., Wich, S. A., Fredriksson, G., Perwitasari-Farajallah, D., Pamungkas, J., & Krützen, M. (2013). Marked population structure and recent migration in the critically endangered Sumatran orangutan (Pongo abelii). Journal of Heredity, 104(1), 2–13.

Nei, M., Tajima, F. & Tateno, Y. Accuracy of estimated phylogenetic trees from molecular data. J Mol Evol 19, 153–170 (1983).

Peakall, R., & Smouse, P. E. (2006). genalex 6: Genetic analysis in Excel. Population genetic software for teaching and research. Molecular Ecology Notes, 6(1), 288–295.

Peakall, R., & Smouse, P. E. (2012). GenAlEx 6.5: Genetic analysis in Excel. Population genetic software for teaching and research—an update. Bioinformatics, 28(19), 2537–2539.

Pocock, R. I. (1939). The fauna of British India, including Ceylon and Burma (Vol. 1, Mammalia, R.B. S. Sewell, Ed.). Taylor and Francis.

Populations, 1.2.30 Copyright (C) 1999, Olivier Langella, CNRS UPR9034

Pritchard, J. K., Stephens, M., & Donnelly, P. (2000). Inference of Population Structure Using Multilocus Genotype Data. Genetics, 155(2), 945–959.

Rousset, F. (2008). genepop ‘007: A complete re-implementation of the genepop software for Windows and Linux. Molecular Ecology Resources, 8(1), 103–106.

Sekar, S., & Karanth, P. (2013). Flying between Sky Islands: The Effect of Naturally Fragmented Habitat on Butterfly Population Structure. PLoS ONE, 8(8), e71573.

Tyagi, A., Singh, S., Mishra, P., Singh, A., Tripathi, A. M., Jena, S. N., & Roy, S. (2016). Genetic diversity and population structure of Arabidopsis thaliana along an altitudinal gradient. AoB PLANTS, 8, plv145.

van Elst, T., Schüßler, D., Rakotondravony, R., Rovanirina, V. S. T., Veillet, A., Hohenlohe, P. A., Ratsimbazafy, J. H., Rasoloarison, R. M., Rasoloharijaona, S., Randrianambinina, B., Ramilison, M. L., Yoder, A. D., Louis, E. E., & Radespiel, U. (2023). Diversification processes in Gerp’s mouse lemur demonstrate the importance of rivers and altitude as biogeographic barriers in Madagascar’s humid rainforests. Ecology and Evolution, 13(7), e10254.

Van Oosterhout, C., Hutchinson, W. F., Wills, D. P. M., & Shipley, P. (2004). micro - checker: Software for identifying and correcting genotyping errors in microsatellite data. Molecular Ecology Notes, 4(3), 535–538.

Wallace, A. R. (1854). On the monkeys of the Amazon. Annals and Magazine of Natural History, 14(84), 451–454.

Wang, W., Qiao, Y., Zheng, Y., & Yao, M. (2016). Isolation of microsatellite loci and reliable genotyping using noninvasive samples of a critically endangered primate, Trachypithecus leucocephalus. Integrative Zoology, 11(4), 250–262.

Wangchuk, T., Inouye, D. W., & Hare, M. P. (2003). A New Subspecies of Golden Langur (Trachypithecus geei) from Bhutan. Folia Primatologica, 74(2), 104–108.

Waples, R. S. (2015). Testing for Hardy–Weinberg Proportions: Have We Lost the Plot? Journal of Heredity, 106(1), 1–19.

