## Supplementary Material for "Population genetics of Himalayan langurs and its taxonomic implications"

**Table S1.** Comparison of genetic diversity estimates across all loci and populations

All loci

| | N | Na | Ho | He |
|---|---|---|---|---|
| **Mean** | 48 | 4.200 | 0.410 | 0.560 |
| **SE** | | 0.371 | 0.045 | 0.040 |

Only including loci in linkage equilibrium

| | N | Na | Ho | He |
|---|---|---|---|---|
| **Mean** | 48 | 4.040 | 0.394 | 0.534 |
| **SE** | | 0.474 | 0.057 | 0.051 |

Abbreviations: $H_O$: Observed heterozygosity, $H_E$: Expected heterozygosity, $N_A$: Mean number of alleles, N: Sample size.

**Figure S1.** Bar plot of Structure

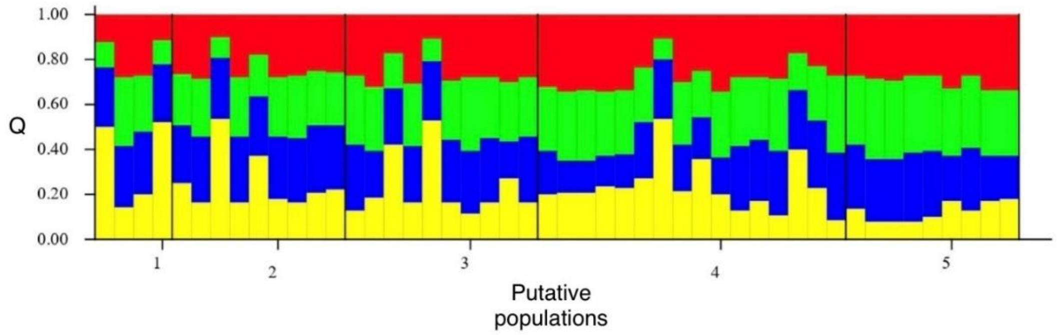


Abbreviations: 1, 2, 3, 4, and 5 correspond to subpopulations of Himalayan langurs separated by river valleys, i.e, nS, S-Y, Y-BG, B-A, eAG respectively.

**Figure S2.** Linear regression of XY plot from Mantel test

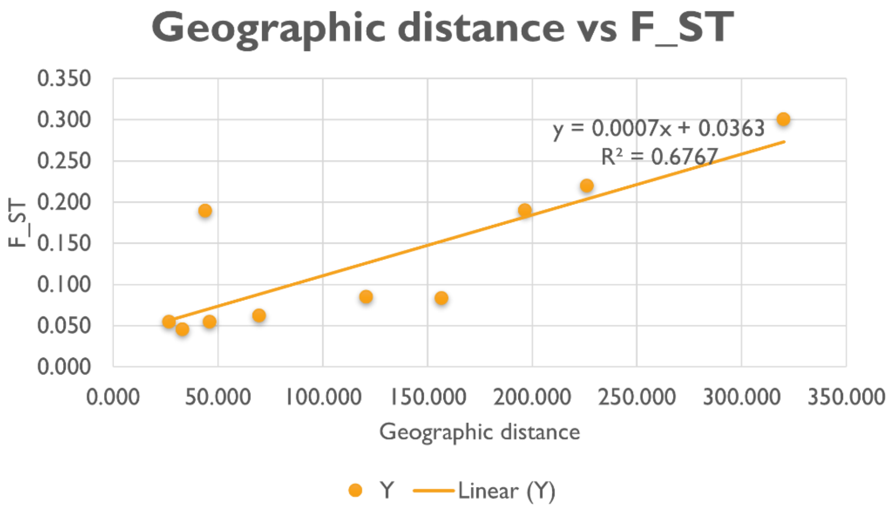
